# Surface tension drives neuronal sorting in magnetically engineered brain-like tissue

**DOI:** 10.1101/2022.12.15.520562

**Authors:** Jose E. Perez, Audric Jan, Catherine Villard, Claire Wilhelm

## Abstract

Engineered 3D brain-like models have advanced the understanding of neurological mechanisms and disease, yet their mechanical signature, while fundamental for brain function, remains understudied. The surface tension for instance controls brain development and is a marker of cell-cell interactions. Here, we engineered 3D magnetic brain-like tissue spheroids composed of intermixed primary glial and neuronal cells at different ratios. Remarkably, the two cell types self-assemble into a functional tissue, with the sorting of the neuronal cells towards the periphery of the spheroids, whereas the glial cells constitute the core. The magnetic fingerprint of the spheroids then allows their deformation when placed under a magnetic field gradient, at a force equivalent to a 70 g increased gravity at the spheroid level. The tissue surface tension and elasticity can be directly inferred from the resulting deformation, revealing a transitional dependence on the glia/neuron ratio, with the surface tension of neuronal tissue being much lower. This provides the underlying mechanical explanation for the exclusion of the neurons towards the outer spheroid region, and depicts the glia/neuron organization as a surface tension-driven sophisticated mechanism that should in turn influence brain development and homeostasis.

## 1. Introduction

Human brain *in vitro* 3D models have since recent years been utilized for recapitulating the properties and physiology of this organ with the purpose of studying brain-penetrating drugs,^[1]^ cell-cell interactions,^[2]^ brain disease^[3]^ and neurological disorders.^[4]^ These models have evolved from 2D organized networks^[5]^ to culture in 3D scaffolds or matrices^[6–8]^, and more recently to complex organoid systems with defined brain regions that more closely approximate the *in vivo* brain cell biology.

The response of brain cells to changes in the tissue’s mechanical properties is known to play important roles in development, physiology, signaling and pathology.^[9]^ It has been evidenced for instance that the stiffness of the cell microenvironment regulates neurite growth^[10]^ and extension, as well as self-renewal and differentiation in neural stem cells.^[11]^ Glial cells may also thrive better on softer substrates, provided they are coupled with suitable adhesion coatings.^[12]^ The tissue surface tension, a biological marker of intercellular binding energy and hence of tissue cohesivity, has been linked to brain gyrification in embryogenesis.^[13]^ The concept of the tissue surface tension arising from the cohesive and adhesive interactions between cell populations and conferring them their liquid-like behavior was introduced by Steinberg many years ago.^[14]^ In brief, it postulates that the liquid-like behavior of intermixed cell populations closely follows that of immiscible liquids, resulting in the sorting out of the cells based on the surface tension of each cell population, as well as the interfacial tension between the phases. Such mechanistic analysis of the behavior of different cell populations as they are interfaced with one another is of particular interest in studies of tissue morphogenesis, with implications in the final purpose of recreating functional 3D tissue models.

In humans, the glia/neuron ratio in terms of cell number sits close to 50:50.^[15]^ However, how this cell arrangement takes place in the case of neuronal and glial cell populations as they become intermixed, as well as the quantification and contribution of each cell population to the overall tissue mechanical properties, have remained largely unexplored despite their relevancy in *in vitro* tissue morphogenesis. Here, we hypothesized that different intermixing ratios of glial and neuronal cells would translate to differential tissue mechanical properties. To investigate this idea, we envisioned the engineering of a brain-like 3D construct that can be physically stimulated in order to retrieve its mechanical properties. A rarely met need in 3D tissue engineering is the controlled organization of single cells via remote manipulation. Magnetic forces are probably first-in-class candidates for this purpose because they can act at a distance.^[16–18]^ Cells can become magnetic through the internalization of biocompatible iron oxide nanoparticles, widely used in cancer theragnostics,^[19]^ drug delivery^[20]^ and already clinically validated as *T_2_* contrast agents in magnetic resonance imaging^[21]^, with recent advances validating their use for *T_1_* contrast.^[22]^ Clinical nanoparticle formulations may additionally exhibit anti-tumor properties.^[23]^

In a self-integrating, all-in-on process, we thus provide here magnetically-formed 3D tissues shaped in less than a day from individual primary glial and neuronal murine cells labeled with biocompatible iron oxide nanoparticles, and with an inherent capacity for further mechanical stimulation throughout their maturation. These active brain-like constructs were used to explore different glia/neuron intermixing ratios to elucidate their distinct contribution to tissue surface tension, elasticity and organization.

## 2. Results

### Spheroid formation after intermixing of neuronal and glial cells

We used a magnetic molding method to form magnetic spheroids after intermixing of primary glial and neuronal cells extracted from mouse embryos (**Figure 1a**). It is based on an initial quick labeling and magnetization step of the cells with iron oxide nanoparticles while in suspension (see Experimental Section). First, a series of spherical molds is made by partially submerging steel round beads of 1 mm in diameter in liquid agarose, held in place by an array of magnets below. After gelation of the agarose, the steel beads are removed with the aid of a magnet, leaving behind a perfectly spherical mold. The magnetized cell populations at the desired intermixed ratio are pipetted into the mold, with the magnet array below acting as an attractor and thus compelling the cells to aggregate into a spherical shape as they fill the mold. The methodology results in spheroids with a high level of sphericity and high compaction that translate to maturation times as short as 1 day.

**Figure 1.**
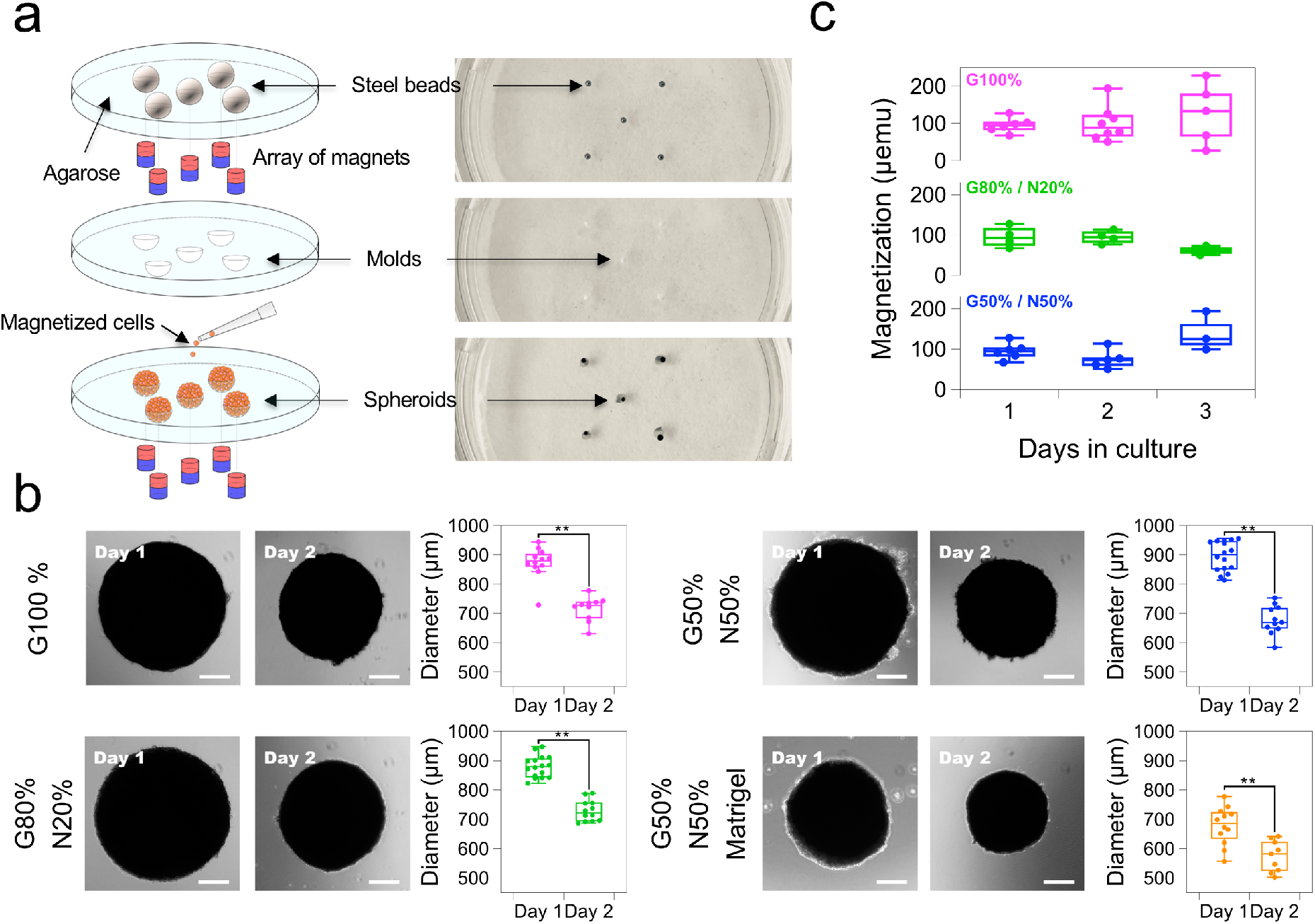
Magnetic molding of neuronal and glial cell spheroids. (a) Spherical molds are produced in agarose using a set of steel beads of 1 mm in diameter and held in place with an array of magnets. Upon removal of the beads, cells that have been pre-labeled with magnetic nanoparticles are attracted into the mold using the same array of magnets in order to obtain spheroid whose shape is defined by the agarose mold. (b) Optical images of spheroid formed after intermixing glial and neuronal cell populations at ratios of G100%, G80% / N20%, G50% / N50% and G50% / N50% plus Matrigel matrix. Quantification of spheroid diameter over a period of 2 days shows a significant decrease in size for all conditions. (c) Spheroid magnetization quantification over 3 days of culture by vibrating sample magnetometry. *n* = 3; \*\**p* < 0.01. Scale bars = 200 μm.

Five different cell ratios were studied in terms of the percentage of cells composing the spheroid, henceforth denoted as glia/neuron (G/N) ratios: G100%, G80% / N20%, G50% / N50%, G20% / N80% and N100%. Additional conditions of cells mixed with Matrigel before the spheroid formation process were produced in order to study a possible effect of the presence of an underlying matrix on the mechanical properties of the spheroids. It should be mentioned that for conditions of G20% / N80% and N100%, the spheroids lost their viability after day 1 of maturation, likely due to them possessing a low threshold cohesivity. We produced fewer spheroids for these conditions.

Typical spheroids for each of the intermixed cell population conditions are shown in Figure 1b. Remarkably, a consistent spheroid shrinkage and compaction can be observed across all the conditions, as quantified by a decrease in spheroid diameter of 15% on average of the original diameter upon spheroid collection at 1 day of maturation in the mold. Of interest too is the fact that for spheroids with Matrigel intermixed the initial diameter after 1 day of maturation was smaller compared to all the non-Matrigel conditions. All formed spheroids possess a magnetic fingerprint bestowed by the labeling of each individual cell with the magnetic nanoparticles, and which forms the basis of the studying of the mechanical properties of this work. Figure 1c shows the evolution of the spheroids’ magnetization over the course of 3 days of maturation, measured by vibrating sample magnetometry at the single spheroid level. The magnetization remains relatively constant over the maturation of the spheroids, in the range of 100 μemu for all conditions, indicating no loss of magnetic properties over this period of time.

Direct observation of the spheroids through fluorescence microscopy revealed the arrangement architecture of the neuronal and glial cell populations within the spheroid. Interestingly, the red glial fibrillary acidic protein (GFAP)-stained glial cells visually appear to arrange themselves towards the core of the spheroid structure. For the G80% / N20% ratio spheroids at day 1 of maturation, neuronal cells appear to cluster in small groups along the periphery of the spheroid, observable by β-tubulin III staining in green (**Figure 2**a). Significant elongation of neuronal processes, presumably axons for the longest ones, is seen inside and in-between these clusters. The development of neuronal branches is remarkably advanced at day 3 of maturation, promoting the connection of initially isolated neuronal clusters (Figure 2b). Increasing the neuronal density to 50% induces the presence of even more neuronal clusters observed at day 1 (Figure 2c). After 3 days, neuronal processes have fully spread at the periphery of the spheroid, with multiple axonal branching and networks appearing to completely cloud over the (red-stained) glial cell population (Figure 2d).

**Figure 2.**
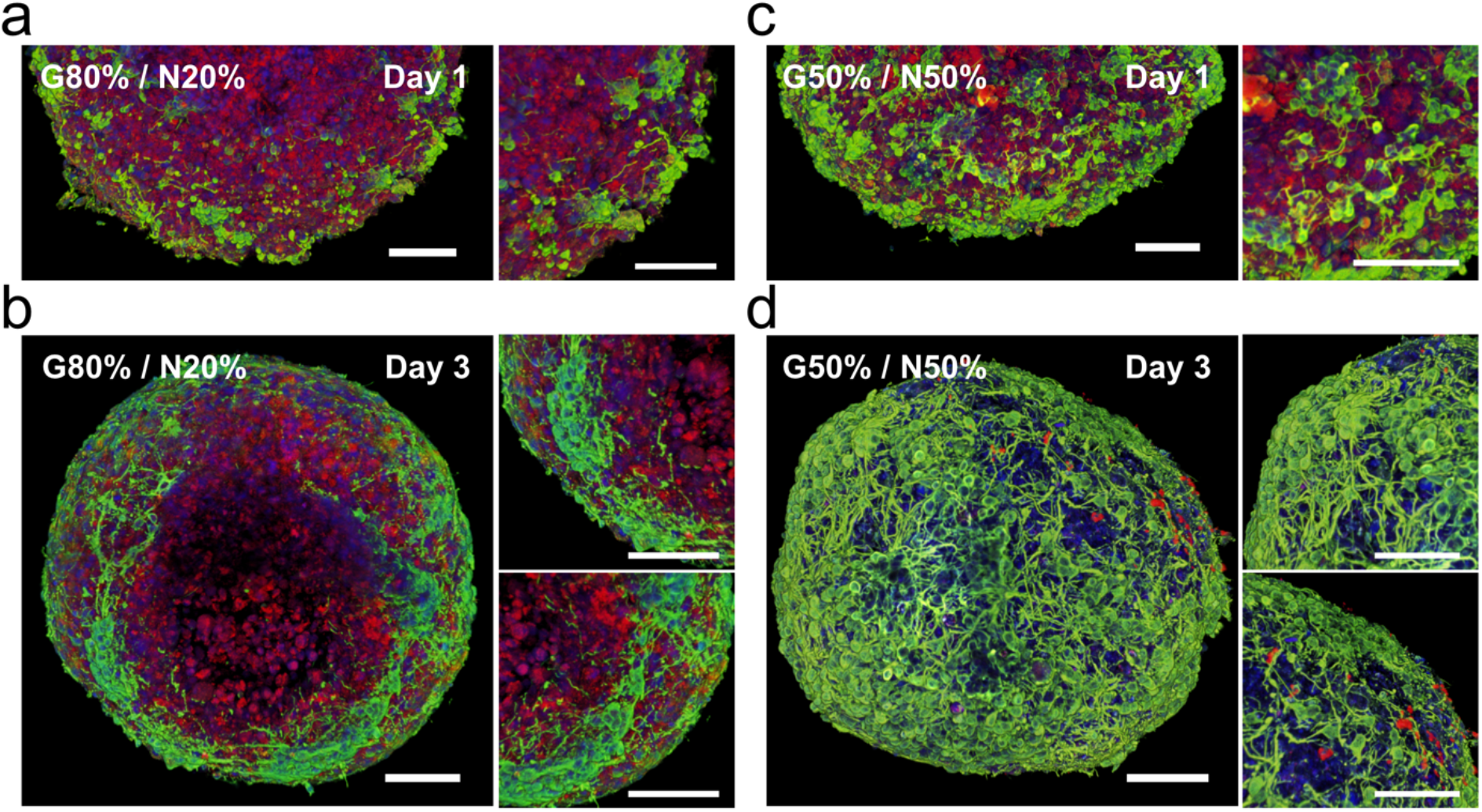
Fluorescence imaging 3D reconstruction of spheroids after glial and neuronal cell intermixing and magnetic molding. (a) and (b) Spheroids at a ratio of G80% / N20% after 1 and 3 days of maturation, respectively. (c) and (d) Spheroids at a ratio of G50% / N50% after 1 and 3 days of maturation, respectively. Fluorescence images show β-tubulin III in green, GFAP in red and DAPI in blue. Scale bars = 100 μm.

It should be noted that the 3D reconstruction of the spheroids shown here was obtained after fluorescence imaging performed with the whole spheroid, representing a visual depth of field of view of approximately 200 μm of tissue due to optical limitations.^[24]^ In spite of this, for all the conditions analyzed, it appears that the intermixing of these two types of cells leads to a specific arrangement where the neuronal cells group themselves in the outer shell of the spheroid, establishing cell-cell communication and axonal networks on top of an inner core of glial cells.

We then proceeded to image the iron oxide nanoparticles organization within the cells after magnetic labeling. For this purpose, thin (20 μm) cross-sections were obtained from the inner central planes of each spheroid (**Figure 3**a). Prussian blue staining of this inner spheroid structure revealed the presence of the blue signal characteristic of iron throughout the whole spheroid, confirming the presence of nanoparticles inside individual cells (Figure 3b). In addition, punctual blue iron signal shows the presence of nanoparticles inside endosomal compartments within the cells. This observation was further confirmed by transmission electron microscopy (TEM) imaging, showing the presence of groups of nanoparticles within cell endosomal compartments (Figure 3c). Overall, the glial cells appeared to internalize the nanoparticles more readily and efficiently, with many of them found densely confined within endosomes. In contrast, for neurons (N100%), the nanoparticles mostly appeared near the cell membrane. Additional images of the nanoparticle localization in glial and neuronal cells are provided in Figures S1 and Figure S2 (Supporting Information), respectively. In the case of neuronal cells, endocytosis events were observed at later spheroid maturation time points (Figure S3, Supporting Information). Thus, after intermixing, glial cells can be systematically identified by the presence of magnetic endosomes (Figure S4 and Figure S5, Supporting Information). Neuronal axonal structures can be observed in the vicinity of glial cells for both of the two conditions of intermixing ratios imaged, with possible synapses being present (Figure S6, Supporting Information). Furthermore, qualitative TEM image analysis also led to the observation that the neuronal cells were more likely to be found at the periphery of the spheroid (Figure S7, Supporting Information).

**Figure 3.**
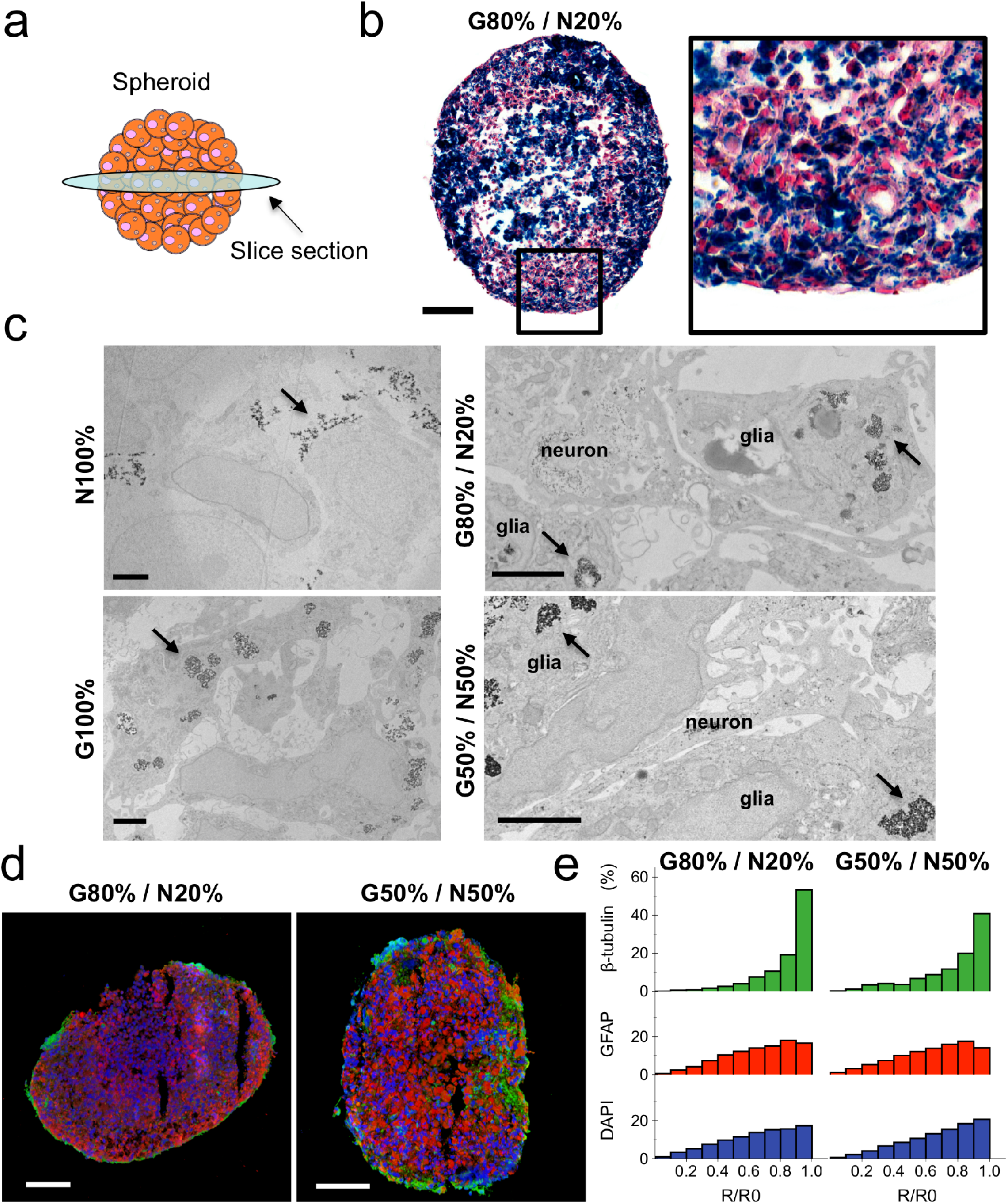
Spheroid cross-section Prussian blue staining imaging, transmission electron microscope imaging analysis and fluorescence imaging and quantification of neuronal and glial cell populations at intermixing ratios of G80% / N20% and G50% / N50%. (a) Spheroid cross-sections were obtained from the center of the spheroid by cryo-sectioning. (b) Prussian blue staining showing the presence of nanoparticles inside the cells, in the form of blue spots suggesting a localization within endosomal compartments. Scale bars = 100 μm. (c) Electron microscopy images showing nanoparticles localized near the cell membrane for the N100% condition, or endosomal compartmentalization for the rest of the conditions, as well as neuronal axonal structures proximal to glial cells. Scale bars = 2 μm. (d) Cross-section fluorescence imaging showing β-tubulin III in green, GFAP in red and DAPI in blue. Scale bars = 100 μm. (e) Radial localization quantification of green-β-tubulin III, red-GFAP and blue-DAPI fluorescent signals according to their position relative to the center of the spheroid. The β-tubulin III signal appears more frequently along the periphery of the spheroid cross-section (R/R0 ≈ 1), whereas the GFAP and DAPI signals are more evenly distributed.

In order to confirm the cell sorting arrangement inferred from 3D reconstruction and TEM observations, we visualized the fluorescent distribution of β-tubulin III, GFAP and cell nuclei (DAPI) in the inner central sections of the spheroids. Results confirmed the localization of the red-stained glial cells mostly in the center of each spheroid plane, whereas the green-stained neuronal cells were observed for the most part at the periphery. This arrangement was remarkably similar for both of the intermixing ratios of G80% / N20% and G50% / N50% (Figure 3d). Additional fluorescence staining images showing this cell arrangement are provided in Figure S8 (Supporting Information). Direct radial localization quantification of the fluorescent intensities of both β-tubulin III, GFAP and DAPI was further assessed to elaborate on this hypothesis. This was done by dividing the spheroid fluorescent planar image using a micrometer-sized grid, and then quantifying the distance between each of the section points within the grid and the center of the spheroid in order to obtain the position of each cell population relative to the spheroid center, given by the ratio of *R/R0*. (see Experimental Section). Results of this analysis evidenced a marked presence of green-β-tubulin III signal along the spheroid periphery (*R/R0* » 1), with red-GFAP and blue-DAPI signals having a more widely distributed position relative to the spheroid center (Figure 3e).

The next step was to exploit the magnetic fingerprint of the spheroids to determine how the different neuronal and glial intermixing ratios would affect the spheroid mechanical properties of surface tension and elasticity. The methodology is shown in **Figure 4**a. It is based on the quantification of the progressive morphological change of the spheroid (i.e. its progressive flattening) when submitted to a magnetic force, until it reaches an equilibrium shape. Briefly, each spheroid is placed on top of a glass coverslip mounted inside an experimentation chamber that permits lateral optical imaging. Then, a magnet of known field gradient is mechanically approached from below the chamber to direct external contact with the glass coverslip (Figure S9, Supporting Information; see Experimental Section). The force then experienced by the spheroid as the magnet approaches thus induces a non-destructive compression deformation along the vertical axis of the spheroid. With the spheroid magnetic moment in the range of 100 μemu (10^-7^ Am^2^) and the gradient generated by the magnet of 170 T m^-1^, the magnetic force (*MvgradB*) acting at the spheroid level is in the range of 15-20 μN. Importantly, it is a volume force of about 5 x 10^4^ N m^-3^, similar to gravity and equivalent to 70 g. Such magnetic-induced gravity is sufficient to flatten the spheroids without damaging the cells, being in the range of what is experienced by cells when under a routine centrifugation step. Figure 4b shows the typical spheroid compression profiles under the applied uniform magnetic field for the glial and neuronal cell intermixing ratios of G100%, G80% / N20% and G50% / N50%. After 5 minutes of magnetic field application the spheroid compression reaches an equilibrium profile, from which the surface tension g can be calculated by solving for Laplace capillary equations. Naturally, a higher spheroid compression, i.e. decrease in overall spheroid height, translates to a lower tissue surface tension. The quantification of the change in spheroid height during the 5 minutes of applied magnetic field is shown in Figure 4c. The G100% spheroids show an overall height loss of 115 μm on average, whereas the G80% / N20% and G50% / N50% ones decrease by 120 and 180 μm on average, respectively. We additionally estimated the spheroid elasticity *E* from the spheroid compression profile at equilibrium following Hertz’s contact theory for an elastic sphere, calculated from the lateral length contact area (2*L*) of the spheroid (Figure 4d), given the known variables of spheroid radius, magnetization, volume and magnetic field gradient (see Experimental Section).

**Figure 4.**
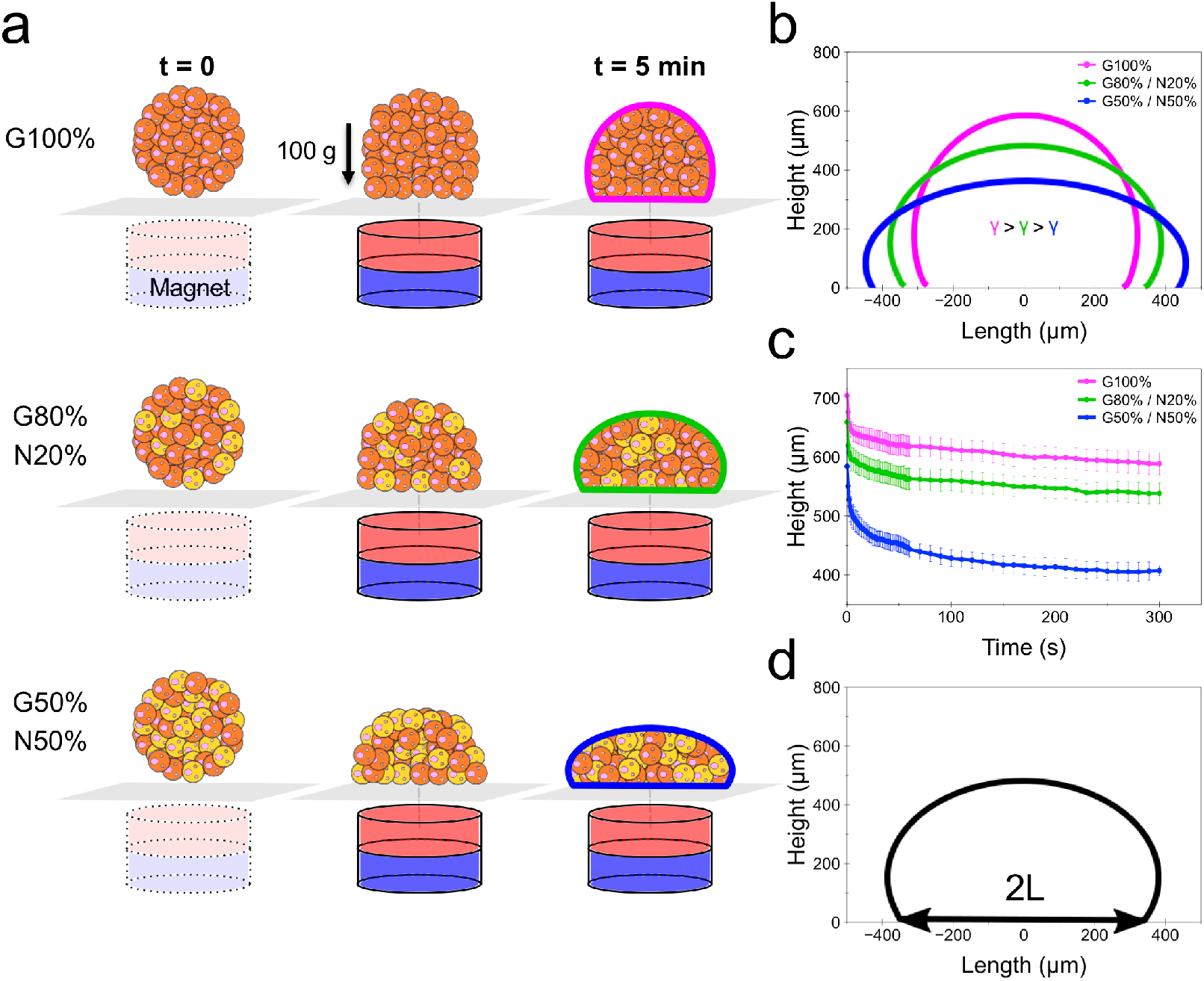
Measurement of the surface tension and elasticity in magnetic spheroids. (a) Diagram showing the followed methodology. A single spheroid is placed on top of a glass coverslip. A magnet is then approached from below in a finely controlled manner and made to be in contact with the glass coverslip, directly below the spheroid position. Upon the applied magnetic field, the spheroid experiences a compression force in the range of 70 g, reaching an equilibrium shape in approximately 5 minutes. (b) Typical spheroid compression profiles for intermixed glial and neuronal cell populations, from which the surface tension g is estimated by solving for Laplace capillary equations. (c) Spheroid height change quantification upon application of the magnetic field over the course of the 5 minutes of compression. Data represent mean ± SEM. (d) Elasticity *E* calculated from the lateral length contact area (2*L*) of the spheroid compression profile at equilibrium, as per Hertz’s contact theory for an elastic sphere.

**Figure 5** demonstrates the dependence of the surface tension and elasticity on the ratio of glial and neuronal cell populations for spheroids at day 1 of maturation. Figure 5a shows the typical optical lateral imaging of the spheroids under magnetic compression for all the conditions tested, along with the spheroids’ final equilibrium profile after 5 minutes of compression. Additional compression profiles are shown in Figure S10 (Supporting Information) It can be visually observed that as the number of glial cells in the spheroid composition increases, the lower the compression is, or conversely, the higher the surface tension of the spheroid. The compression effect is mitigated by the addition of Matrigel into the mixture, more remarkably so for the spheroids with fewer neuronal cells. The quantification of the surface tension g and the elasticity *E* thus revealed that both of these biomechanical parameters are proportional to the percentage of glial cells in the spheroid composition (Figure 5b), with the median surface tension g for the N100%, G20% / N80%, G50% / N50%, G80% / N20% and G100% equal to 4, 15, 17, 37 and 35 mN m^-1^, respectively. The median elasticity *E* was found to be at 60, 114, 280, 593 and 857 Pa for the respective spheroid composition conditions. The addition of Matrigel into the cell mixture significantly increased the surface tension and elasticity for all conditions.

**Figure 5.**
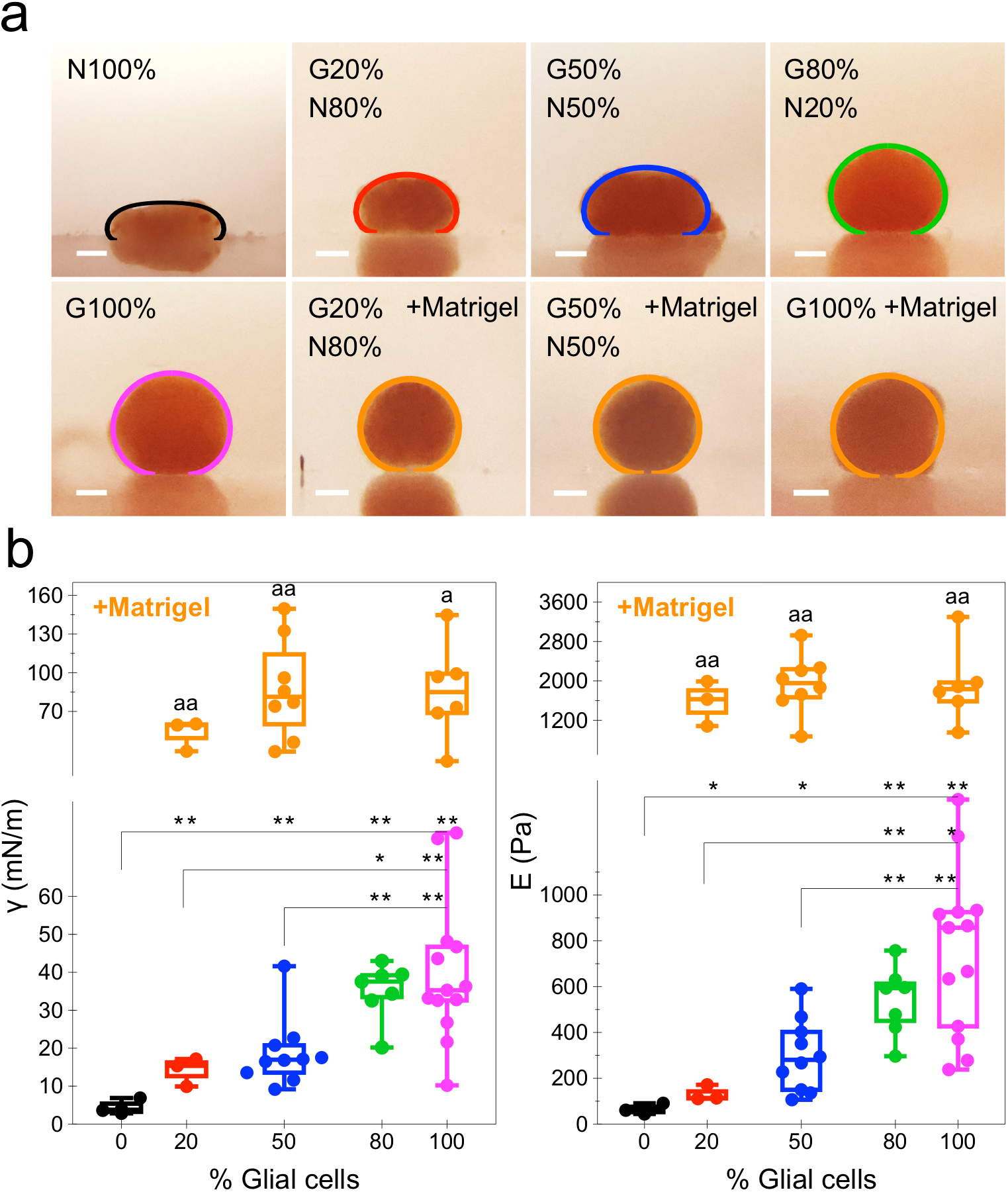
Spheroid surface tension and elasticity dependence on the glia/neuron cell population ratio. (a) Lateral optical imaging of spheroids for all measured cell intermixing ratio conditions with the corresponding equilibrium shape fit obtained after 5 minutes of compression under the applied magnetic field. (b) Spheroid surface tension and elasticity quantification for spheroids at day 1 of maturation and at intermixing ratios of N100%, G20% / N80%, G50% / N50%, G80% / N20% and G100%, as well as the G50% / N50% and G100% ratio spheroids pre-mixed with Matrigel matrix. *n* = 3, except for groups where glial cells ≤ G20%, where *n* = 2; \**p* < 0.05, \*\**p* < 0.01; ^a^*p* < 0.05, ^aa^*p* < 0.01 vs. same cell ratio condition without Matrigel. Scale bars = 200 μm.

Next, we extended this analysis to further spheroid maturation stages, with similar results. For the conditions analyzed, the spheroids’ final equilibrium profile after 3 days showed an overall increase in height, an indication of an increase in tissue cohesivity (**Figure 6**a). Naturally, such a change in the equilibrium profile translates to an increase in surface tension, and a decrease in the contact area 2*L*, hence a higher elasticity. Indeed, the surface tension showed a median increase for all the ratio conditions (Figure 6b). For the G50% / N50% spheroids, the surface tension increased from 17 to 29 mN m^-1^ after 2 days of maturation, albeit it being found to be not significant. On the other hand, spheroids at G80% / N20% showed a significant increase after day 3, from 37 up to 71 mN m^-1^. The neuron-less condition G100% increased from 35 up to 64 and 107 mN m^-1^ for day 2 and 3 of maturation, respectively. The elasticity parameter saw similar results, with overall significant increases compared with the initial day 1 of maturation, with G50% / N50% spheroids reaching a maximum elasticity of 702 Pa, whereas the G80% / N20% and G100% reached values of 1438 and 1585 Pa at day 3, respectively. The addition of Matrigel further increased both biomechanical parameters, although the changes were only significant at day 1, and at day 2 only for the G50% / N50% spheroid condition (Figure 6c).

**Figure 6.**
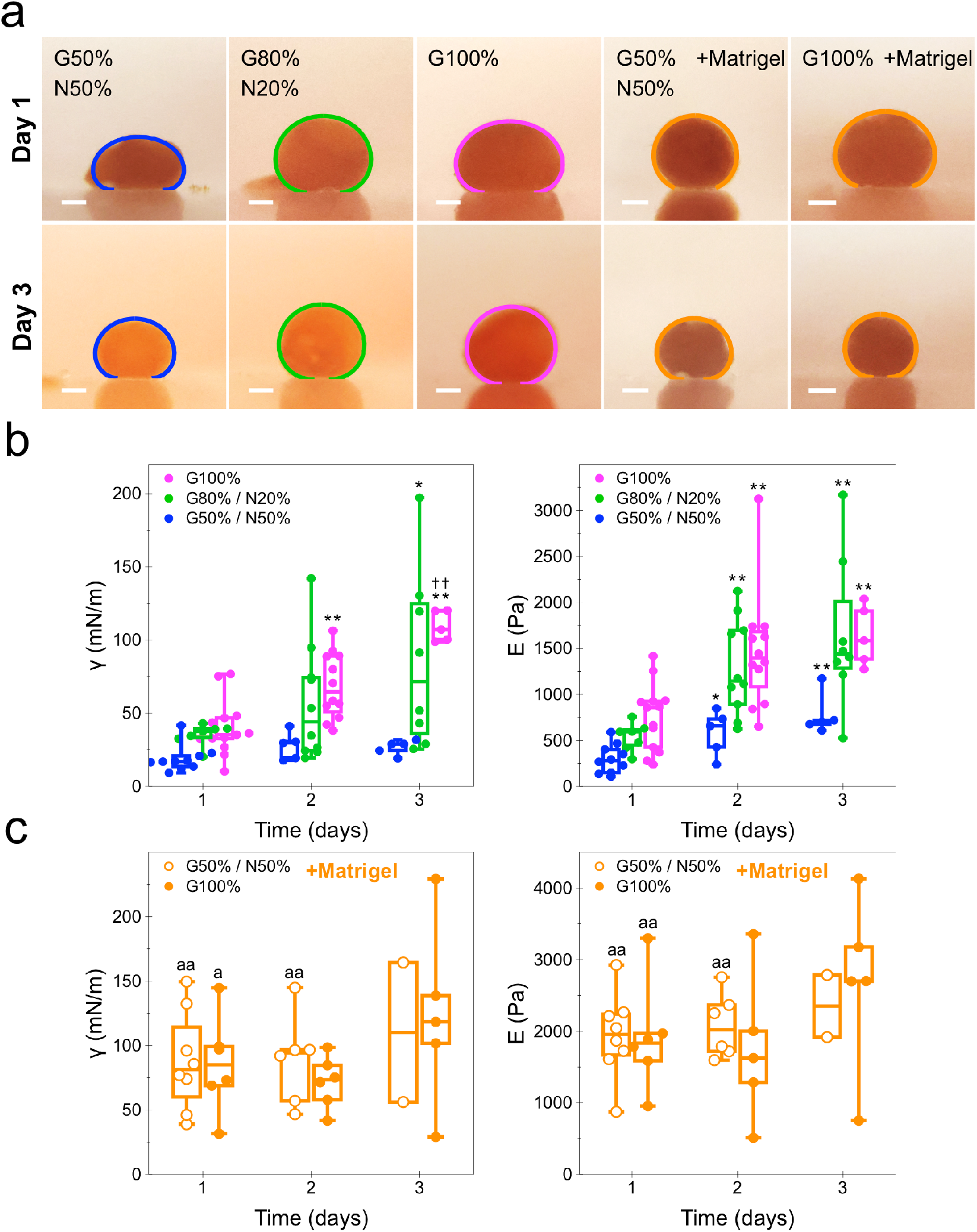
Spheroid surface tension and elasticity dependence on the glia/neuron cell population ratio over 3 days of maturation. (a) Lateral optical imaging of spheroids for all measured cell intermixing ratio conditions with the corresponding equilibrium shape fit obtained after 5 minutes of compression. Images show spheroids at day 1 and day 3 of maturation. (b) Spheroid surface tension and elasticity quantification for spheroids at these maturation time points and for intermixing ratios of G50% / N50%, G80% / N20% and G100%, as well as the G50% / N50% and G100% ratio spheroids premixed with Matrigel matrix. *n* = 3; *n* = 2 for later maturation time point spheroids with Matrigel; *\*p* < 0.05, \*\**p* < 0.01 vs. day 1; ^††^*p* < 0.01 vs. day 2; ^a^*p* < 0.05, ^aa^*p* < 0.01 vs. same cell ratio condition without Matrigel, at same day of maturation. Scale bars = 200 μm.

## 3. Discussion

We validated an all-in-one magnetic based strategy for brain-like tissue formation, maturation and mechanical stimulation. We used magnetically-labeled primary glial and neuronal cells intermixed at different ratios. Both cell types show the capacity of nanoparticle uptake, yet glial cells appear to more readily internalize and endocytose them. The nanoparticle uptake process renders the endosomes magnetic, conferring in turn magnetism to each cell. It is thanks to this cell magnetization process that they can be aggregated under an external magnetic field within minutes, rapidly adopting a spherical shape in the process. Mature spheroids are obtained in less than 1 day, permitting the direct observation of important early stage tissue morphogenesis processes that are otherwise unattainable with typical spheroid formation techniques. Indeed, the arrangement of the cells on the outer layers of the spheroid changes drastically between the first and third days of maturation, as neuronal cells extend their axonal network and spread over the glial cell population, which remain mostly confined in the center of the spheroid. Neurite length may reach about 100 μm after day 1, which exceeds the length observed in 2D cultures in astrocyte conditioned medium on Matrigel.^[25]^ This highlights the fact that our culture conditions promote healthy, fast growing neurons.

The idea of the envelopment of one cell population over another as they become intermixed was initially described by the differential adhesion hypothesis, which states that the surface tension, itself a result of the cohesive and adhesive forces between the component cells, bestows the tissue its liquid-like behavior.^[26]^ It is therefore the surface tension, or the intensity of cohesion among the cells, that drives the envelopment of one cell population over another. Thus, the cell population with the lower surface tension tends to spread over or envelop the one with the higher surface tension.^[27]^ Here, we show that primary neuronal cells sort themselves around glial cells after cell intermixing and magnetic aggregation into a spheroid, with this specific architecture being maintained over a period of 3 days of maturation as the neuronal axonal network spreads and extends along the spheroid periphery.

In order to further prove this concept, we sought out to elucidate the specific contribution of the glial and neuronal cells to the overall spheroid surface tension. This biomechanical property is obtained by subjecting the spheroid tissue to a controlled force and allowing it to reach an equilibrium shape,^[28]^ which is reached as the applied force is balanced out by the tissue cohesive forces driving its spherical shape conformation. Here, the surface tension was measured using a novel technique that exploits the magnetic fingerprint of the spheroid due to the labeling of cells with magnetic nanoparticles: by subjecting the spheroid to an external field gradient, it experiences a deformation along its vertical axis and reaches the equilibrium shape within minutes. The surface tension measurement of the glial and neuronal spheroids at different intermixing ratios directly confirmed the previously observed cell sorting behavior, with results indicating that the increase in spheroid surface tension is proportional to the number of glial cells in the mixture. On the other hand, the presence of neuronal cells tends to slow down the time-dependent increase of g. Thus, the lower surface tension of the neuronal cells is the main driver of their spreading and arrangement over the glial cells in the early stages of the spheroid formation process (< day 1). Although there exist scarce or no reports in the literature of the surface tension of primary brain cells, this parameter has been studied in glioblastoma aggregate models, with the surface tension found in the range of 7 - 46 mN m^-1^ depending on the tumoral cell line,^[29,30]^ within the same range of our reported value of 35 mN m^-1^ for the G100% spheroids. The addition of Matrigel significantly increased the surface tension for the G20 / N80% and G50% / N50% ratio conditions compared to the spheroids without matrix, at 3.7- and 4.6-fold at day 1 of maturation. This effect is explained by an increase in spheroid cohesivity by the addition of a matrix. The surface tension increase due to the addition of Matrigel in the G100% spheroids was overall lower, at 2.4-fold at day 1. This suggests that the surface tension in glial tissue alone, being the highest between all the measured conditions, possesses a threshold cohesivity that cannot be further increased by external inputs.

We further took advantage of the equilibrium shape reached after magnetic compression of the spheroids to infer their elastic properties. For the G50% / N50% ratio condition, which more closely resembles the brain composition, the elasticity was found to be around 280 - 700 Pa depending on the spheroid maturation time. These values are in agreement with previously reported elastic moduli in spheroids with a similar composition, at 100 – 200 Pa,^[31]^ and that of bulk murine hippocampal tissue, found to be in the 300 – 660 Pa range by scanning^[32]^ and atomic force microscopy.^[33]^ Interestingly, it is at this specific substrate elastic modulus range where neuronal differentiation and β-tubulin III expression both peak.^[11]^ The addition of Matrigel into the cell mixture increased spheroid elasticity by 11.8-fold in G20% / N80%, 6.5-fold in G50% / N50% spheroids and 2.5-fold in G100% spheroids at day 1. This is expected, given that Matrigel alone possesses an elastic modulus of approximately 450 Pa.^[34]^ Our elasticity measurements confirm that, at the fundamental level of neuronal and glial cells intermixing, the brain-like tissue spheroids remain relatively soft in comparison with other types of tissues. Indeed, both these types of cells have been evidenced to be relatively soft when measured individually, with glial cells found to be softer than neurons.^[32]^ In contrast to this, we measured a higher elasticity for pure glial spheroids. This suggests that the mechanical properties at the spheroid tissue level do not reflect the properties of individual cells. Indeed, astrocytes are capable of expressing fibronectin^[35]^ and laminin,^[36]^ which may induce a reinforcement of adhesive contacts with neurons and lead to a higher elastic modulus.

## 4. Conclusion

To the best of our knowledge, this is the first report of a viable reconstitution of spheroids from intermixed primary glial and neuronal cells after a rapid magnetic aggregation step. The resulting magnetically active brain-like tissue spheroids can thus be controlled by remote magnets. We used this magnetic fingerprint to retrieve the biomechanical properties of surface tension and elasticity from a magnetic-driven deformation. Remarkably, we evidenced a transition from very low surface tension to much higher ones by the increase of the proportion of glial cells in the neuron-glia mix. Not only this is the first determination of the important difference in the individual contribution of glial and neuronal cells to the overall tissue surface tension, but it additionally mirrors the impressive local organization observed at the spheroid level, with neurons being sorted out to the spheroid periphery. The 3D brain tissue model recapitulates some of the brain physiology, and provides insight into the predetermination of cell population sorting and arrangement in the brain imposed by the surface tension, recalling tissue embryogenesis.

## 5. Experimental Section

### In vivo extraction of neuronal and glial cells

Cortical neurons and glial cells were harvested from the cortical hemispheres dissected from C57BL/6 mouse embryo brains at E15-16 (Charles River Laboratories). Pregnant mice were euthanized by cervical dislocation, and the embryos were surgically removed and euthanized by a spinal cord incision at the neck level before brain extraction. The procedure was carried out three independent times, each one approximately yielding 5-6 embryo brains. After extraction of both cortical hemispheres, they were rinsed thoroughly using Gey’s Balanced Salt Solution (GBSS, Thermofisher), and then dissociated in Dulbecco’s Modified Eagle’s medium (DMEM, Thermofisher) containing 25 mg mL^-1^ of papain (Sigma Aldrich) during 10 minutes at 37 °C. The enzymatic dissociation was halted by the addition of 10% fetal bovine serum (FBS, Thermofisher). Samples were then dissociated in DMEM medium supplemented with DNase I (Sigma Aldrich) with the aid of a pipette. The harvested cells were then centrifuged at 80 g for 6 minutes and resuspended in DMEM medium supplemented with 5% FBS, 1% N-2 (Thermofisher), 2% B-27 (Thermofisher) 2 mM GlutaMAX^®^ (Thermofisher), 1 mM sodium pyruvate (Thermofisher) and 20 μg mL^-1^ gentamicin (Thermofisher), henceforth denoted as DMEMc. All experimental handling of neuronal cells was performed in suspension, right after collection and processing.

For glial cell extraction and culture, the cortical hemispheres were rinsed in GBSS and then transferred to a petri dish previously coated with 15 μg L^-1^ of Poly-L-ornithine (Sigma Aldrich). Each cortex was then cut into small pieces in Modified Eagle’s Medium (Thermofisher) supplemented with 10% FBS, 2mM GlutaMAX^®^, 1 mM sodium pyruvate and 20 μg mL^-1^ gentamicin. Tissue pieces were mechanically dissociated by vigorous pipetting in the culture dish, and the obtained cell suspension was incubated for one week at 37 °C in a humidified incubator at 5% CO_2_, changing the culture medium every 3 days. Before magnetic labeling, the glial cells were detached with 1X TrypLE™ Express Enzyme (Thermofisher) after a 10 minute incubation at 37 °C.

### Magnetic labeling of cells

The magnetic nanoparticles used for cell labeling were produced using the standard procedure by co-precipitation of iron salts. Briefly, FeCl_3_ (3 g) and FeCl_2_ (8.1 g) were diluted in 5 mL deionized (DI) water with 25% ammonium hydroxide used as a precipitating agent (pH 10). After 20 minutes of heating at 90°C, the solution was decanted on magnet, washed with acetone and H_2_O, centrifuged at 8000 rpm for 10 minutes and re-dispersed in 10 mL of DI water. Citrate (2 g in 20 mL DI water) was added, and the resulting solution was heated at 90°C for 1 hour under vigorous agitation, resulting in both nanoparticle oxidation and citrate chelation to ensure colloidal stability. The nanoparticle diameter was 8 ± 1.7 nm, as estimated by TEM.

For magnetic labeling, the neuronal and glial cell populations were both counted and intermixed in DMEM medium (without serum) at the corresponding ratios. Mixed populations were then centrifuged at 100 g for 5 minutes and resuspended in serum-free RPMI medium (Gibco^®^) with magnetic nanoparticles at [Fe] = 50 mM and supplemented with citrate at 5 mM. The cells were magnetically labeled in this solution during 5 minutes at 37 °C in a humidified incubator at 5% CO_2_ and while under shaking motion.

### Magnetic molding of spheroids

The spheroids were formed using the magnetic molding method^[37]^. Briefly, 2% agarose in phosphate buffered saline (PBS) is used to create molds in a petri dish. The molds’ spherical shape is defined by immersing 1 mm steel beads in the liquid agarose and holding them in place using an array of cylindrical neodymium magnets (Supermagnete). The steel beads are removed after agarose gelation and the molds are then sterilized by UV exposition for 30 minutes and filled with PBS. Immediately after labeling, the cells were centrifuged at 100 g for five minutes and each condition of mixed neuronal and glial cells was then resuspended in 50 μL of DMEMc, mixing thoroughly with the aid of a pipette to break out aggregates of cells. Using the same array of neodymium magnets placed below the dish, and hence the molds, 7 μL of the magnetic cell suspension (approximately 100,000 cells) were pipetted into the spherical molds, being thus magnetically attracted towards the direction of the mold and filling it up. The PBS is aspirated from the dishes and replaced with DMEMc, and the formed spheroids embedded in the molds are left to mature overnight at 37°C in a humidified incubator at 5% CO_2_. The next day, the spheroids were collected with the aid of a cut 1 mL pipette tip and subsequently incubated in Neurobasal Medium (Thermofisher) supplemented with 1% FBS, 2% B-27, 1% L-glutamine (Thermofisher) and 1% sodium pyruvate.

For the formation of spheroids containing Matrigel^®^ (Corning^®^) matrix, the initial cell population was mixed with approximately 40% of Matrigel solution in volume before pipetting into the molds. All Matrigel and cell handling procedures were performed at 4 °C to avoid gelation of the matrix. Approximately 14 μL of the cell/matrix suspension were slowly pipetted into the molds.

### Spheroid magnetization quantification

The magnetization of individual spheroids was quantified using a vibrating sample magnetometer (Quantum Design, Versalab), with each measuremenent performed at 300 K over a range of 0 – 3 T. Data shown refers to bulk magnetization expressed in μemu.

### Immunofluorescence staining

At the desired maturation time point, spheroids were fixed in 4% paraformaldehyde for 1 hour at room temperature. They were then washed in PBS and permeabilized in 0.1% Triton X-100 in PBS for 15 minutes at room temperature, and then subsequently blocked in a solution consisting of 2% bovine serum albumin solution in PBS with 0.1% Triton X-100 for 3 hours at room temperature. Spheroids were then incubated in anti-GFAP primary antibody (G4546, Sigma Aldrich) at a 1:200 dilution in blocking solution at 4 °C for a 48 hours period, followed by incubation in anti-β-tubulin isotype III primary antibody (T5076, Sigma Aldrich) at a 1:500 dilution in blocking solution at 4 °C overnight. The spheroids were then washed thoroughly in PBS and incubated in Alexa Fluor 594^®^-conjugated goat anti-rabbit IgG secondary antibody (A11037, Thermofisher) at a 1:500 dilution in blocking solution at 4 °C overnight. After washing in PBS, spheroids were incubated in Alexa Fluor 488^®^-conjugated goat anti-mouse IgG secondary antibody (A11029, Thermofisher) at a 1:400 dilution in blocking solution for 2 hours at room temperature. Lastly, after washing with PBS, spheroids were incubated in a 300 nM DAPI solution (D3571, Invitrogen) at a 1:500 dilution in PBS at 4 °C overnight. A spheroid z-stack section of approximately 200 μm was imaged using a Leica DMi8 inverted confocal microscope (Leica Microsystems) using a 25x water immersion objective.

### Spheroid cryosectioning and Prussian blue staining

After fixation in paraformaldehyde, cryosectioning of the spheroids and Prussian blue staining were carried out by the Plateforme Histologie, Immunomarquage, Microdissection laser (HistIM, Institut Cochin, Paris, France). The obtained cryosections possessed a thickness of 20 μm. Immunofluorescence staining and imaging of spheroid cryosections were performed as previously described.

### Image analysis and fluorescence quantification

Image analysis and 3D spheroid reconstruction were done using the ImageJ open software. For the analysis of the localized radial fluorescent intensity of the neuronal and glial cell populations, each image cross-section was divided in 40 x 40 μm sections. Each of the section points was then assigned an x,y coordinate. Using a custom Python code, each set of coordinates was then individually compared to the x,y coordinates of the spheroid center (manually set) in order to obtain the *R/R0* ratio, where *R* denotes the distance from each section point to the x,y coordinates of the spheroid center, and with *R0* denoting the radius of the spheroid from its center to the closest point of the periphery along the analyzed x,y coordinates of the section point. Thus, the *R/R0* ratio directly measures how far each data point is from the spheroid center. This analysis was performed for each section of the grid comprising the complete spheroid in the image, and for all three fluorescence channels imaged (green for β-tubulin III, red for GFAP and blue for DAPI).

### Transmission electron microscope imaging

Spheroids were fixed in 2% glutaraldehyde in 0.1 M cacodylate buffer (Sigma Aldrich) for 1 hour at room temperature. Individual spheroids were then contrasted with 0.5% oolong tea extract and post-fixed in 1% osmium tetroxide (Sigma Aldrich) and 1.5% potassium cyanoferrate (Sigma Aldrich). Samples were then gradually dehydrated in graded ethanol, at steps from 30% to 100% ethanol, and subsequently embedded in Epon resin. Thin slices of around 70 nm were imaged using a Hitachi HT 7700 transmission electron microscope operated at 80 kV.

### Surface tension and elasticity quantification

In order to obtain the surface tension measurement at the spheroid level, spheroids were placed inside a chamber filled with phenol-free cell medium and possessing two microscope glass slides on its sides to permit imaging, as well as an additional glass coverslip at the bottom of the chamber. A cylindrical 6 x 6 mm neodymium magnet (Supermagnete), generating a uniform magnetic field gradient of *gradB* = 170 T m^-1^ within a cylindric volume of 2 mm in height and 2 mm in diameter, was then approached with fine precision and made to come in contact with the glass coverslip at the bottom of the chamber using a vertical lift stage (X-VSR20A, Zaber Technologies Inc.). The full system was placed inside a custom-built thermoregulated chamber supplemented with CO_2_ (Microscope Heaters - Digital Pixel Limited, Brighton, United Kingdom). As each spheroid possesses a magnetic fingerprint conferred by the labeling with magnetic nanoparticles, they experience a compression deformation in the direction of the magnetic field gradient as the magnet approaches and touches the glass coverslip beneath. The spheroids’ lateral compression profiles were monitored live with a Canon EOS R6 digital camera coupled with a Canon MP-E 65 mm f/2.8 1-5x Macro lens until equilibrium compression shape was reached (approximately 5 minutes). The surface tension parameter can then be calculated from the spheroid shape in equilibrium when compressed by integrating the Laplace law of capillarity^[38]^ to extract the capillary constant *c*:

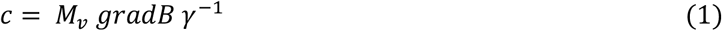

where *M_v_* is the volume saturation magnetization of the spheroid, previously calculated through vibrating sample magnetometry, *gradB* the magnetic field gradient applied and g the macroscopic surface tension of the spheroid.

The spheroid elasticity quantification was done following Hertz’s contact theory for an elastic sphere, estimated by equation (2):

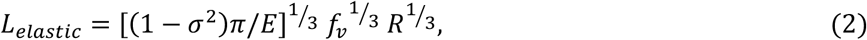

where 2*L* is the lateral length contact area of the spheroid when compressed, *σ* is the Poisson ratio, here set to ½, *f_v_* the volumetric force exerted on the spheroid by the magnetic field gradient, equal to *M_v_ gradB, R* the initial radius of the spheroid and *E* its elastic modulus.

### Statistical analysis

Statistical analysis was done using the Student’s two-Sample t-test, with statistical significance considered for values of *p* < 0.01 or *p* < 0.05.

## Supporting information

Supporting information is available from the Wiley Online Library or from the authors.

## Conflict of interest

The authors declare no conflict of interest.

## Ethics approval statement

The experimental use of laboratory mice was performed in accordance with the European Community guidelines of animal care, under the Institut Curie license #C75-05-18, 24/04/2012, reporting to “Comité d’Éthique en matière d’expérimentation animale Paris Centre et Sud” (National registration #59).

## Acknowledgements

The authors would like to thank Gaëtan Mary for help with the analysis of the fluorescent microscopy images, Christine Péchoux at INRAe for TEM processing and analysis, Yoann Lalatonne for help with the synthesis of magnetic nanoparticles, the Plateforme Mesures Physiques à Basses Températures (Sorbonne University) for the magnetometry analysis, and the HistIM platform at Institut Cochin for spheroid cryo-sectioning and staining.

We also thank the Institut Pierre-Gilles de Gennes (IPGG) with Equipex “Investissements d’avenir” program ANR-10-IDEX-0001-02 PSL and ANR-10-LABX-31-34.

This work was supported by the ANR grant ANR-19-CE09-0029 and the European Union grant ERC-CoG MaTissE 648779.

## Data availability statement

Datasets utilized in the study are available from the authors upon reasonable request.

## Table of contents entry

Neurons self-sort towards the periphery of magnetic brain-like tissue spheroids after intermixing with glia, driven by their overall lower surface tension. Indeed, the magnetically reconstituted spheroids show a transition from low to high surface tension that is dependent on the proportion of glia, which provide most of the tissue cohesive signature.

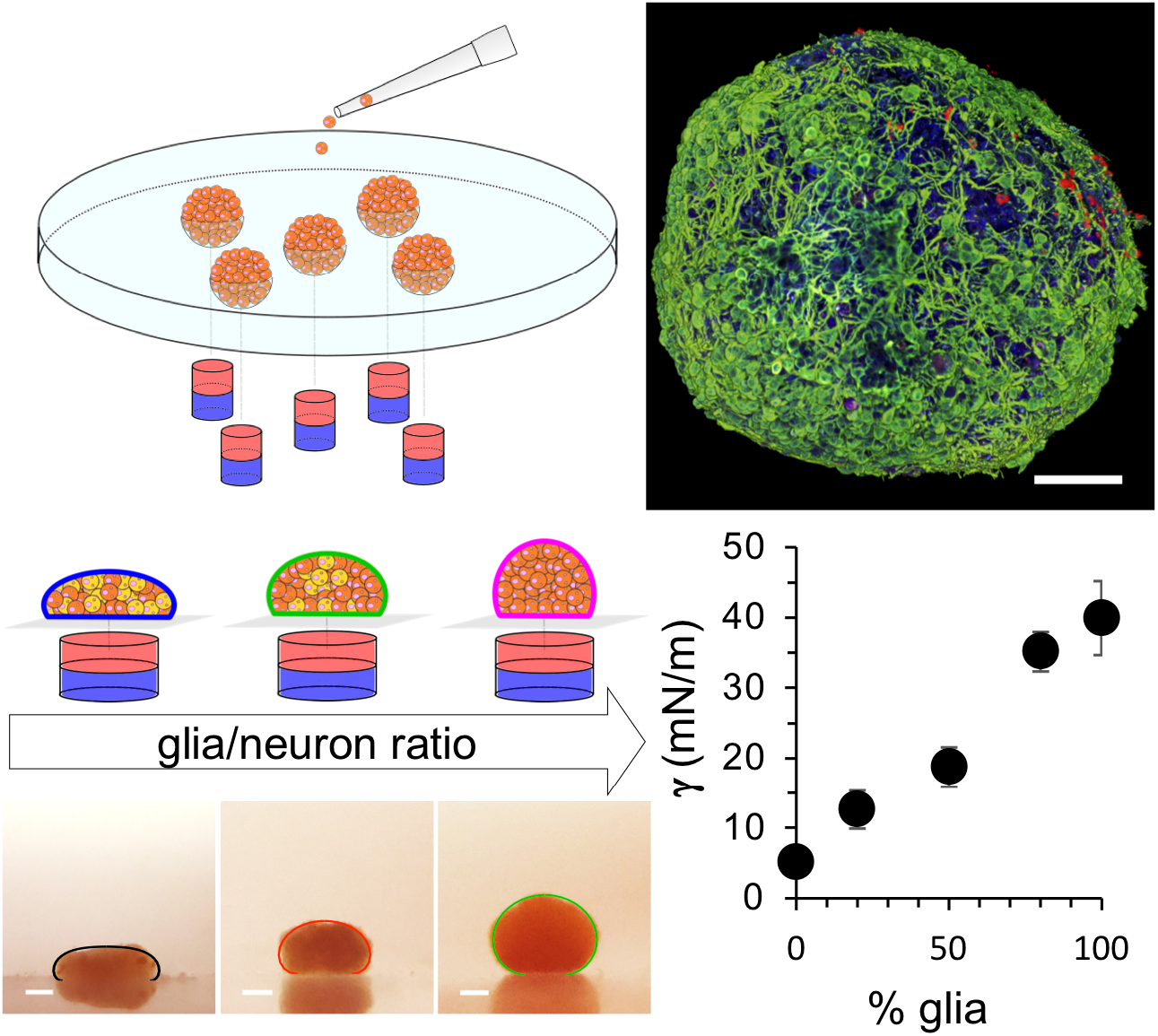

## Notes

### Competing Interest Statement

The authors have declared no competing interest.

