## Supporting Information for "Surface tension drives neuronal sorting in magnetically engineered brain-like tissue"

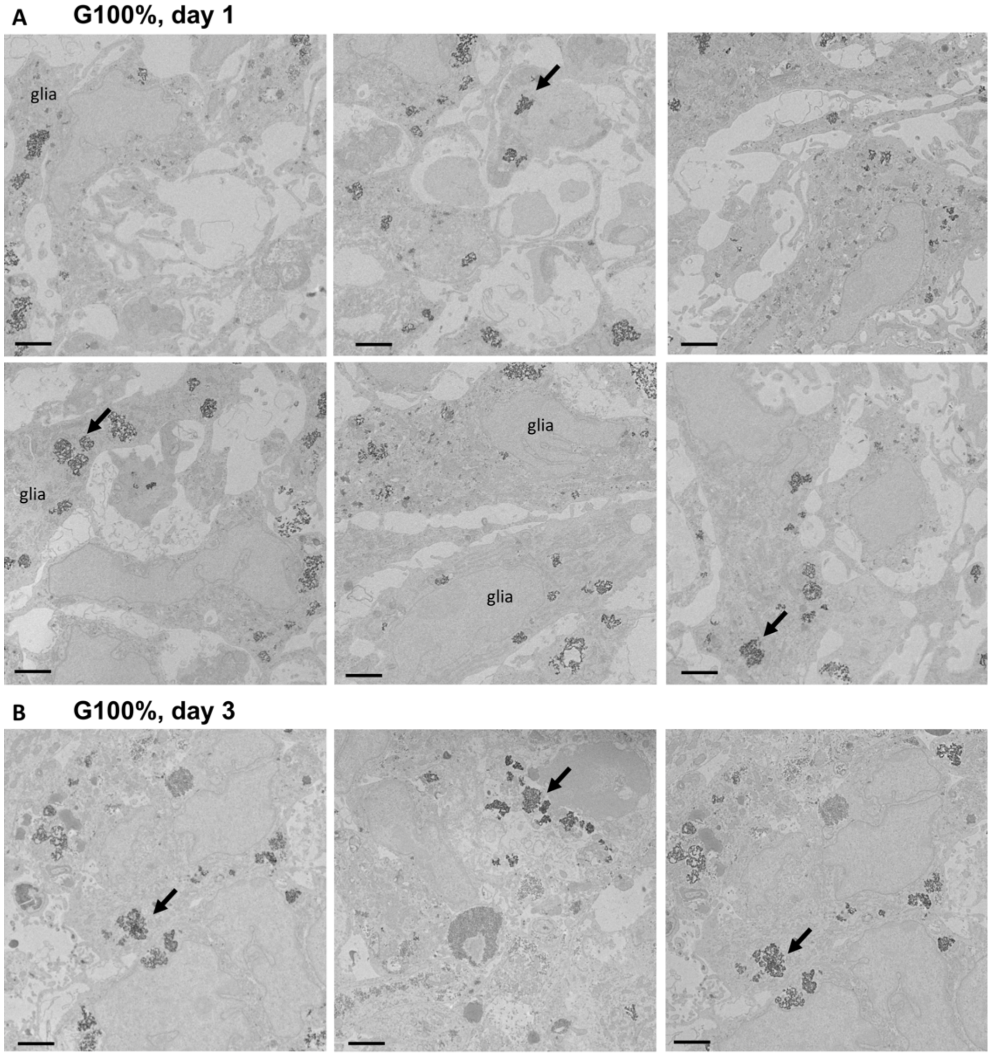


**Figure 1**. TEM images showing endosomal compartmentalization of the nanoparticles in G100% spheroids at (A) day 1 and (B) day 3 of maturation. Arrows indicate instances of nanoparticles confined within endosomes. Scale bars = 2 μm.

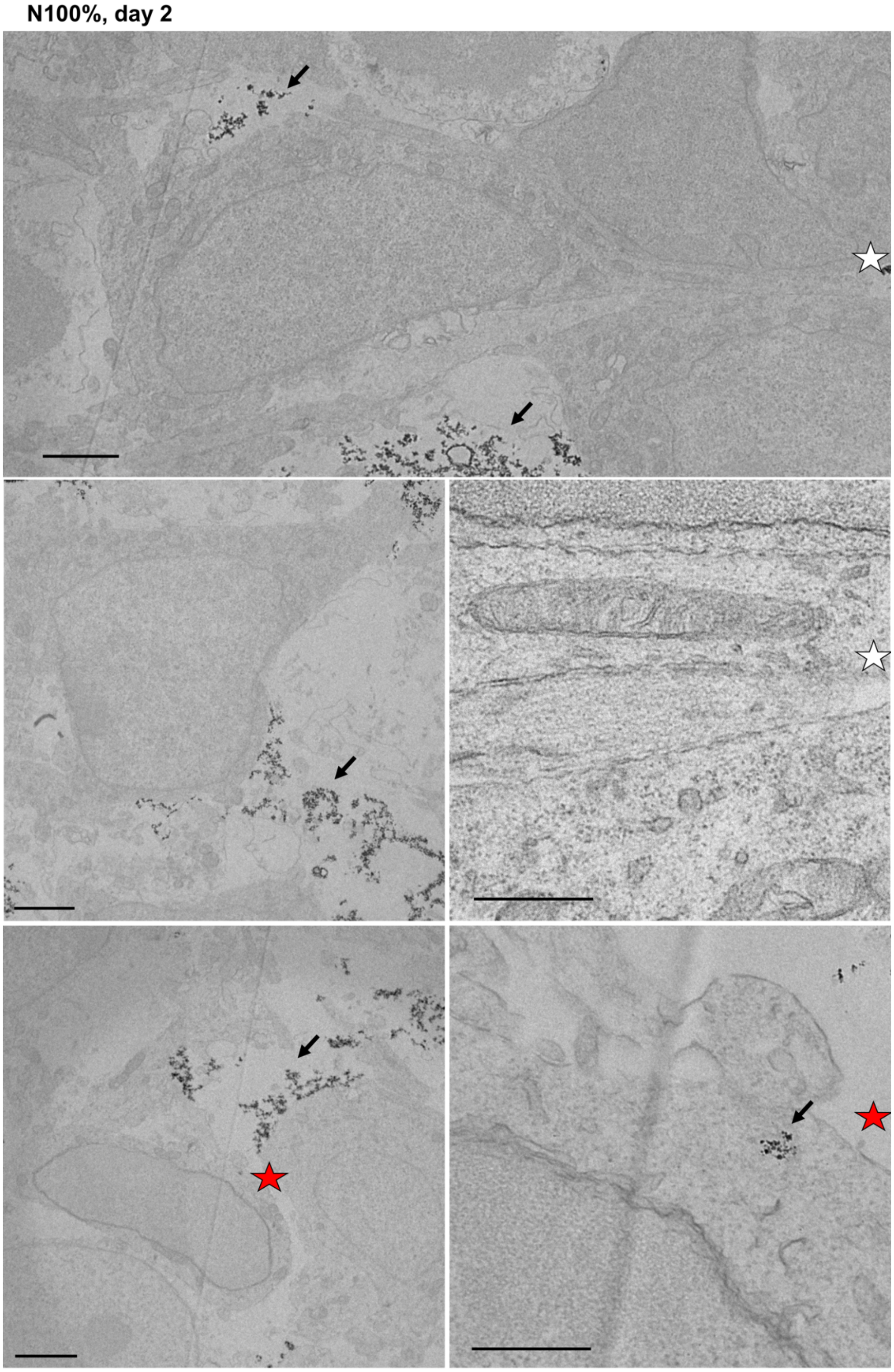


**Figure 2**. TEM images showing the localization of nanoparticles near the cell membrane or at early endosomes (black arrows) in N100% spheroids at day 2 of maturation. Stars indicate concomitant magnification areas. Scale bars = 2 µm.

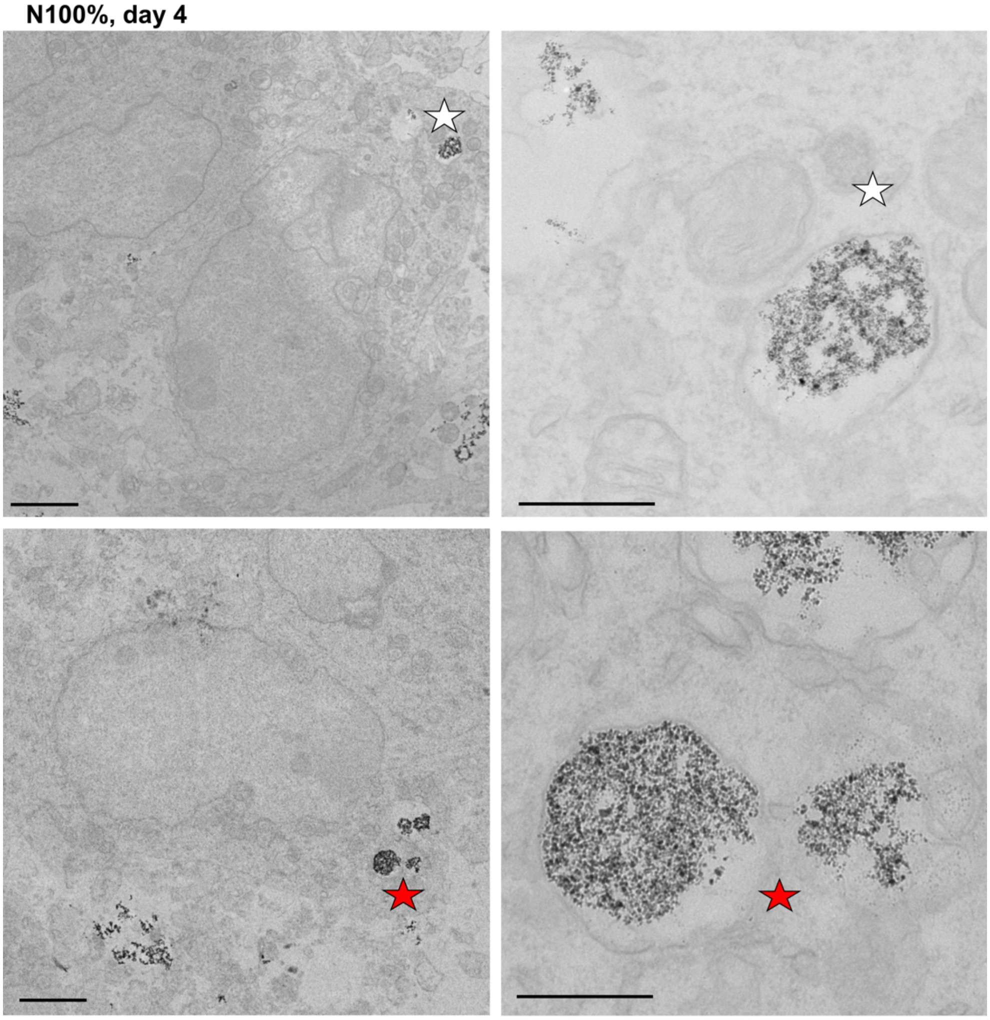


**Figure 3**. TEM images showing the localization of nanoparticles within endosomal compartments in N100% spheroids after 4 days of culture. Stars indicate concomitant magnification areas. Scale bars = 2 µm.

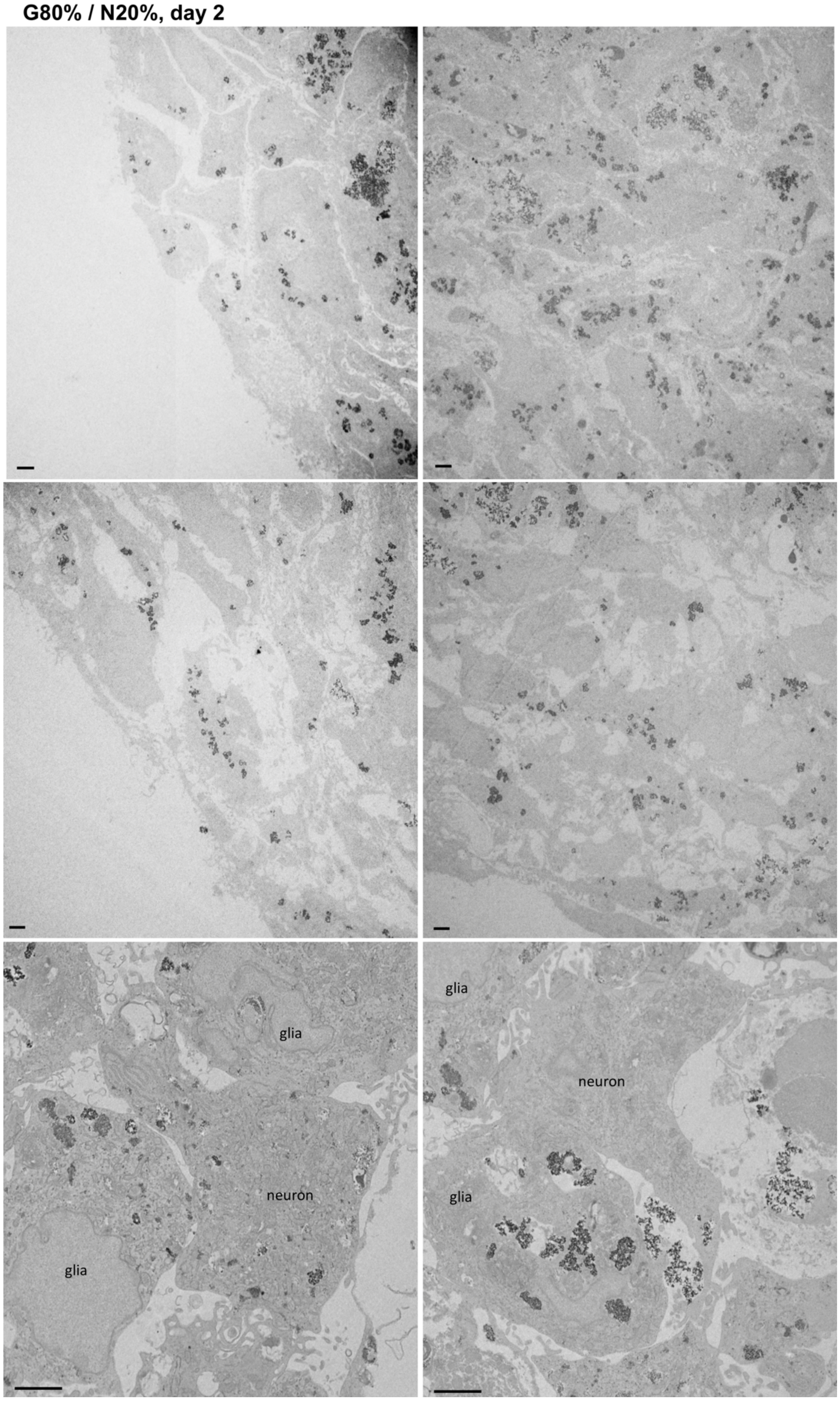


**Figure 4**. TEM images showing the confinement of nanoparticles within endosomes in spheroids formed after intermixing cells at a ratio of G80% / N20%. The glial cell population can be identified by the presence of magnetic endosomes deep within the cellular structure. Scale bars = 2 µm.

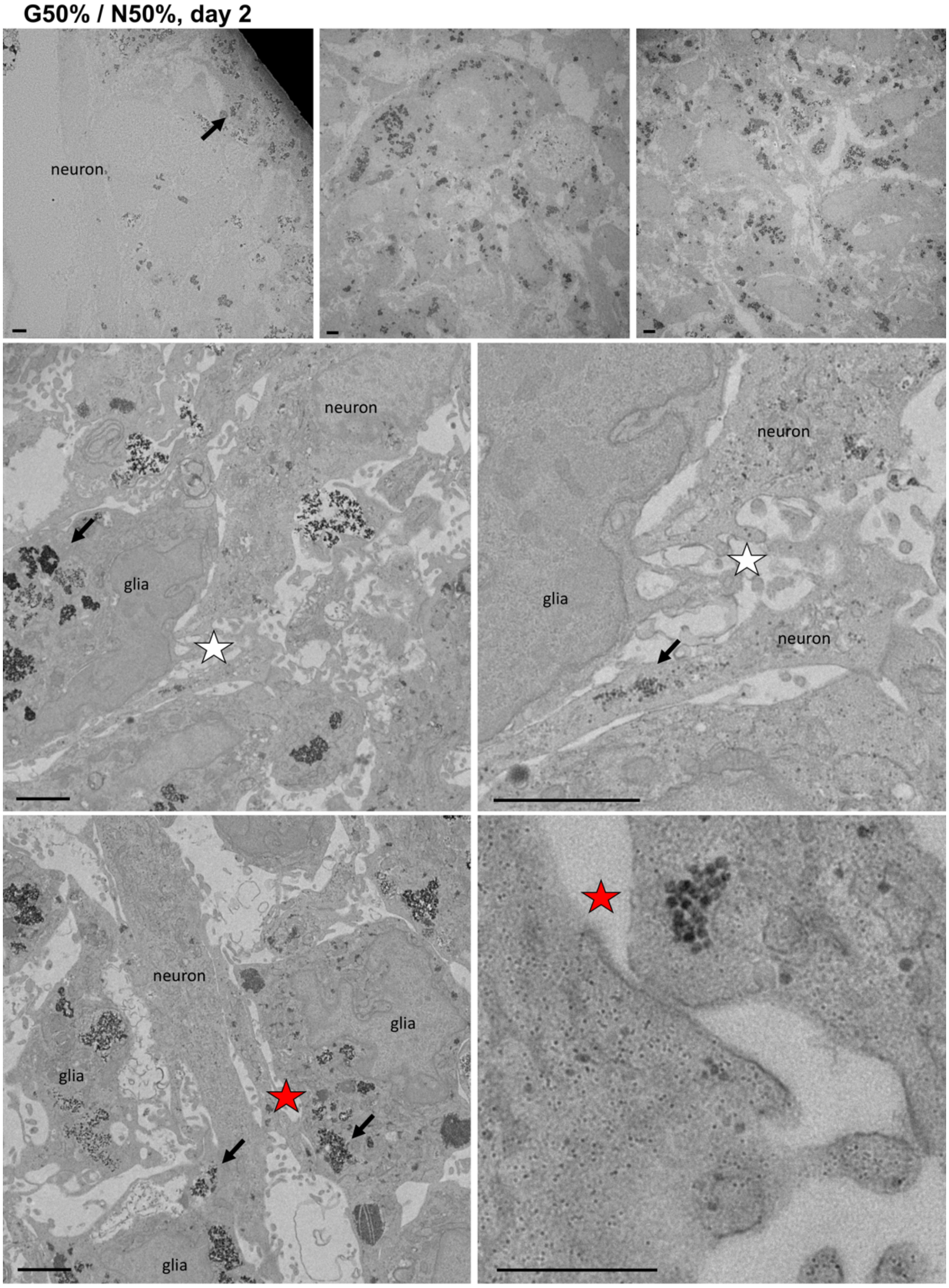


**Figure 5**. TEM images showing the localization of nanoparticles in spheroids formed after intermixing cells at a ratio of G50% / N50%. Arrows denote the presence of nanoparticles near the cell membrane in the case of neuronal cells, or in endosomal compartments for glial cells. Stars indicate concomitant magnification areas. Neuronal structures can be seen in contact with glial cells in the magnified sections. Scale bars = 2 µm.

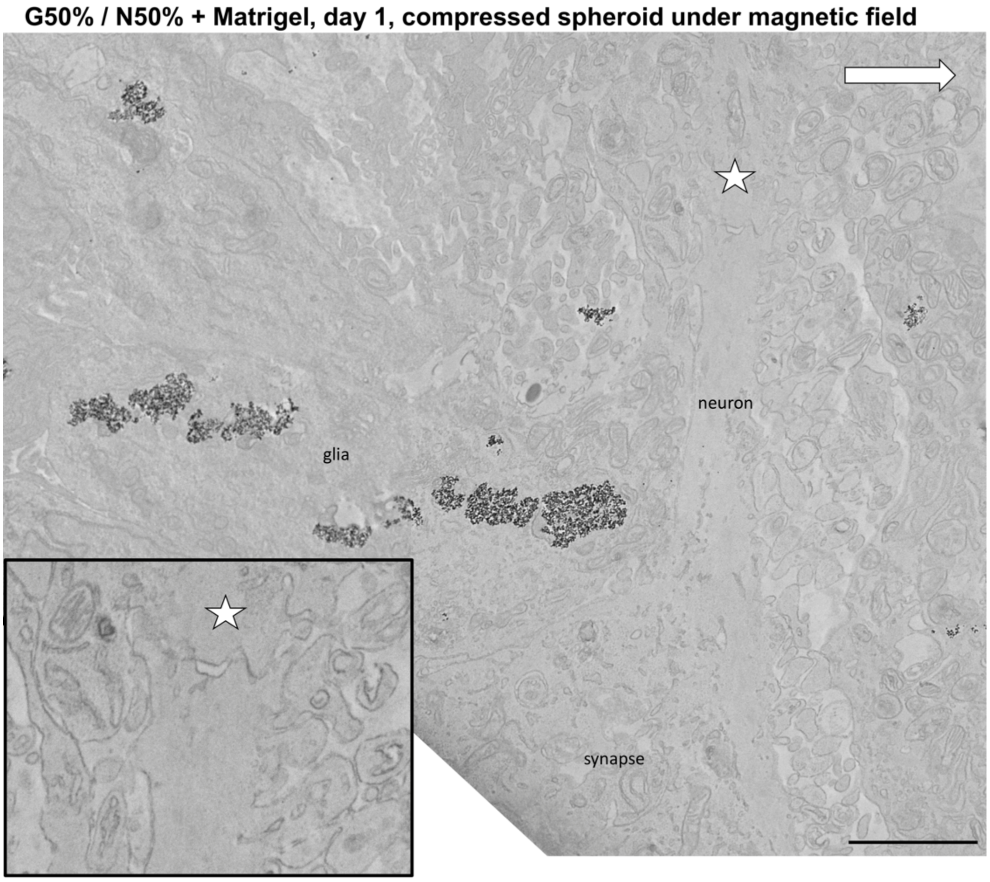


**Figure 6**. TEM image showing a synapse appearing to be located at the end of a neuronal axonal body. The cross-section of the spheroid was obtained while in a compressed state after application of the magnetic field gradient (direction of the gradient is indicated by a white arrow). Stars indicate concomitant magnification areas. Scale bars = 2 µm.

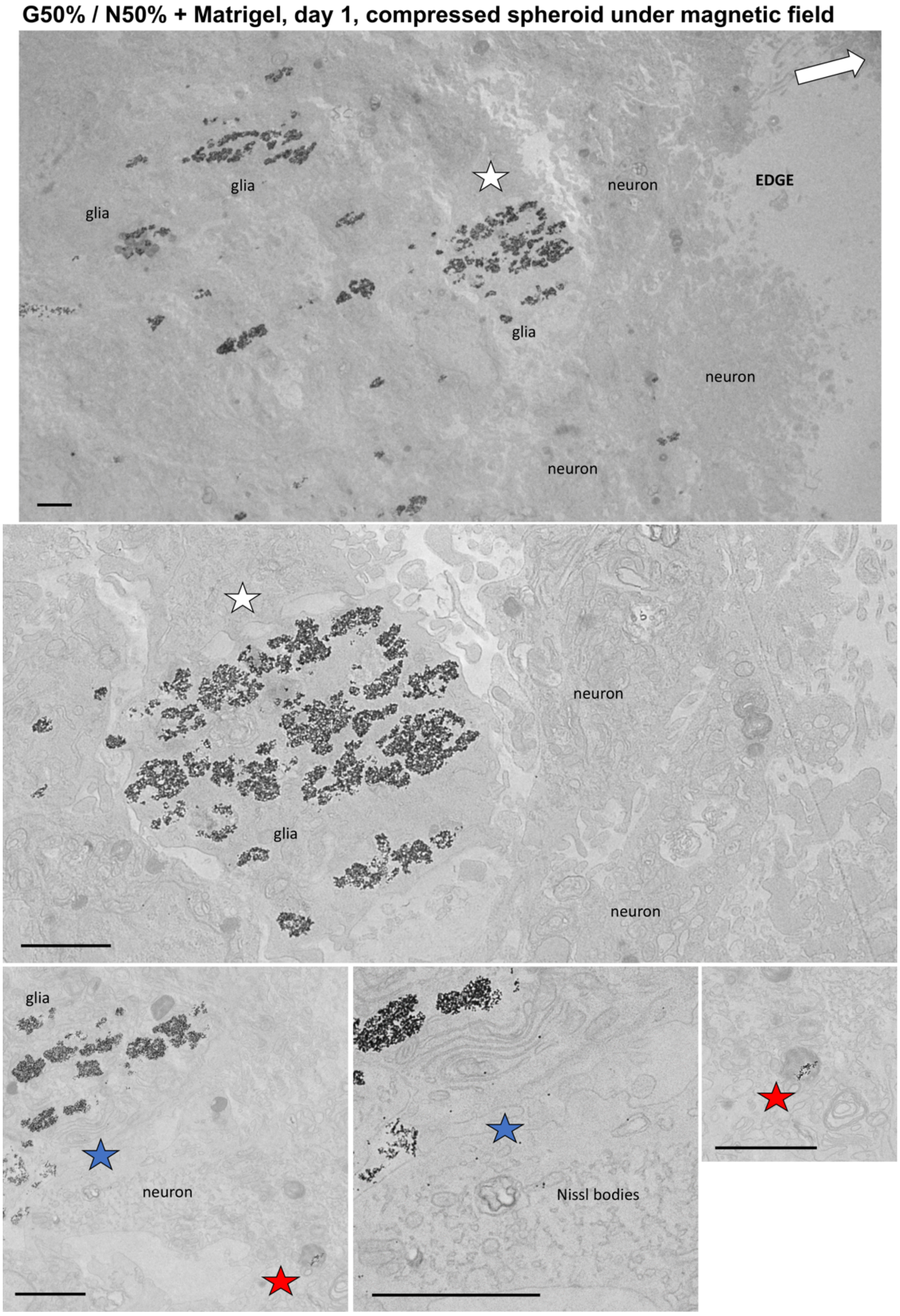


**Figure 7**. TEM images showing the spheroid cross-section localization of the glial and neuronal cell populations after intermixing at a ratio of G50% / N50% with Matrigel matrix. Cross-section of the spheroid was obtained while in a compressed state after application of the magnetic field gradient (direction of the gradient is indicated by a white arrow). Magnetic endosomes can be observed in alignment with the direction of the magnetic field gradient. Neuronal cells locate mostly at the edges of the tissue section. Nissl bodies can be observed in a neuronal cell (lower panel). Stars indicate concomitant magnification areas. Scale bars = 2 µm.

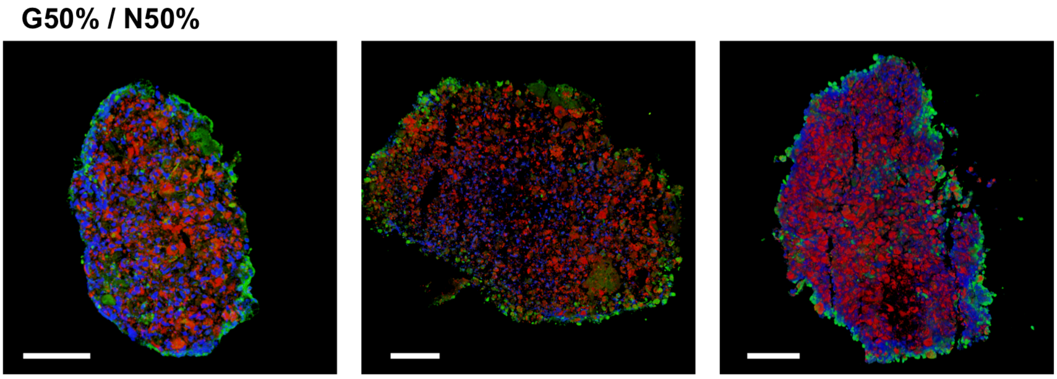


**Figure 8**. Fluorescence imaging of cross-sections of spheroids of neuronal and glial cells at an intermixing ratio of G50% / N50%. Staining shows β-tubulin III in green, GFAP in red and DAPI in blue. The green β-tubulin III signal appears more prominently at the spheroid periphery. Scale bars = 100 µm.

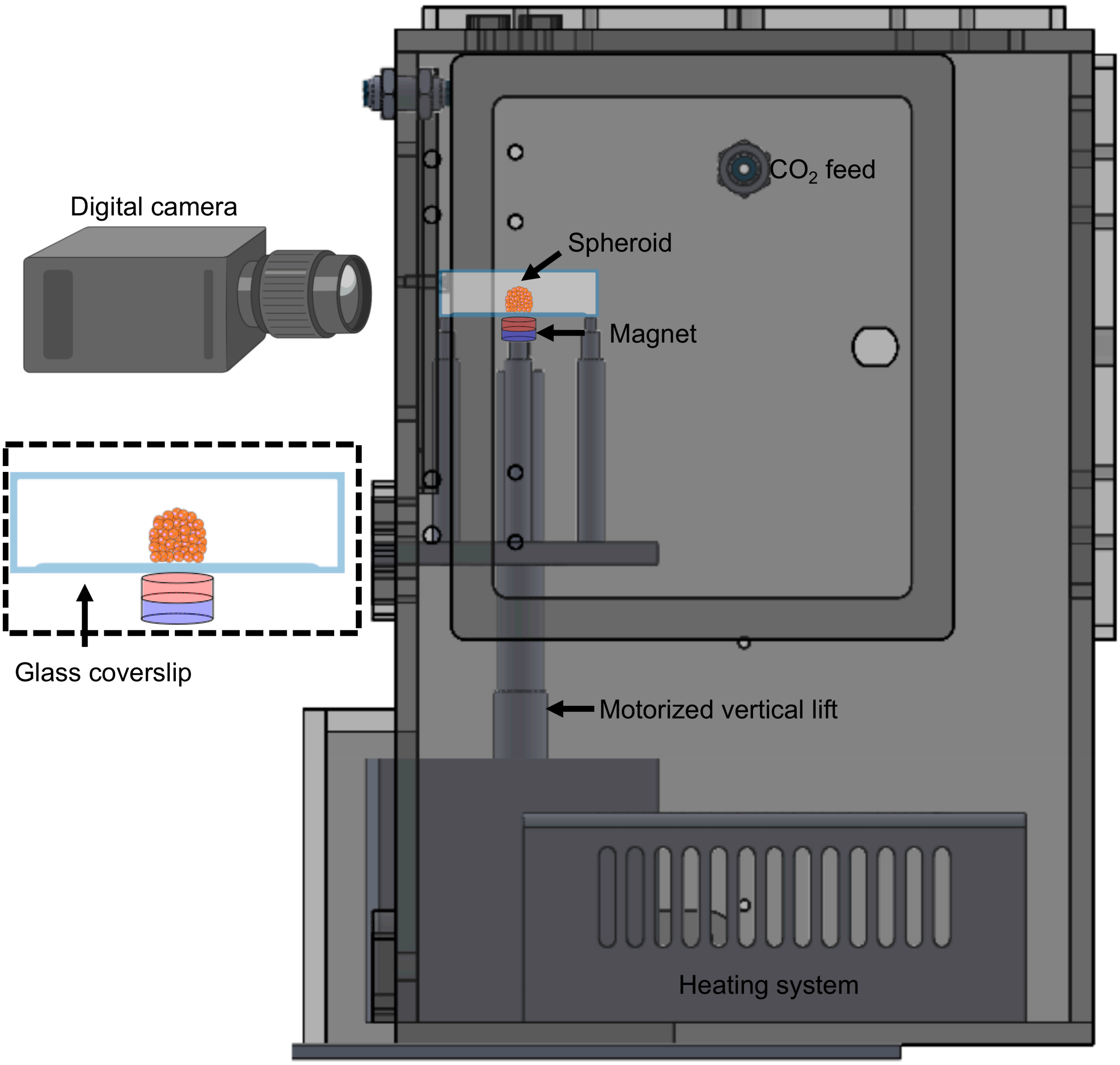


**Figure 9**. Thermoregulated chamber system used for spheroid compression. The custom-built chamber (Microscope Heaters - Digital Pixel Limited, Brighton, United Kingdom) provides a temperature regulation heating system and a feed for $CO_2$. A X-VSR20A vertical lift stage (Zaber Technologies, Inc.) was mounted inside the chamber in order to mechanically tune and control the approach of the magnet below the experimentation chamber (direct external contact with the glass coverslip). A Canon EOS R6 digital camera coupled with a Canon MP-E 65 mm f/2.8 1-5x Macro lens was used to image the spheroid compression.

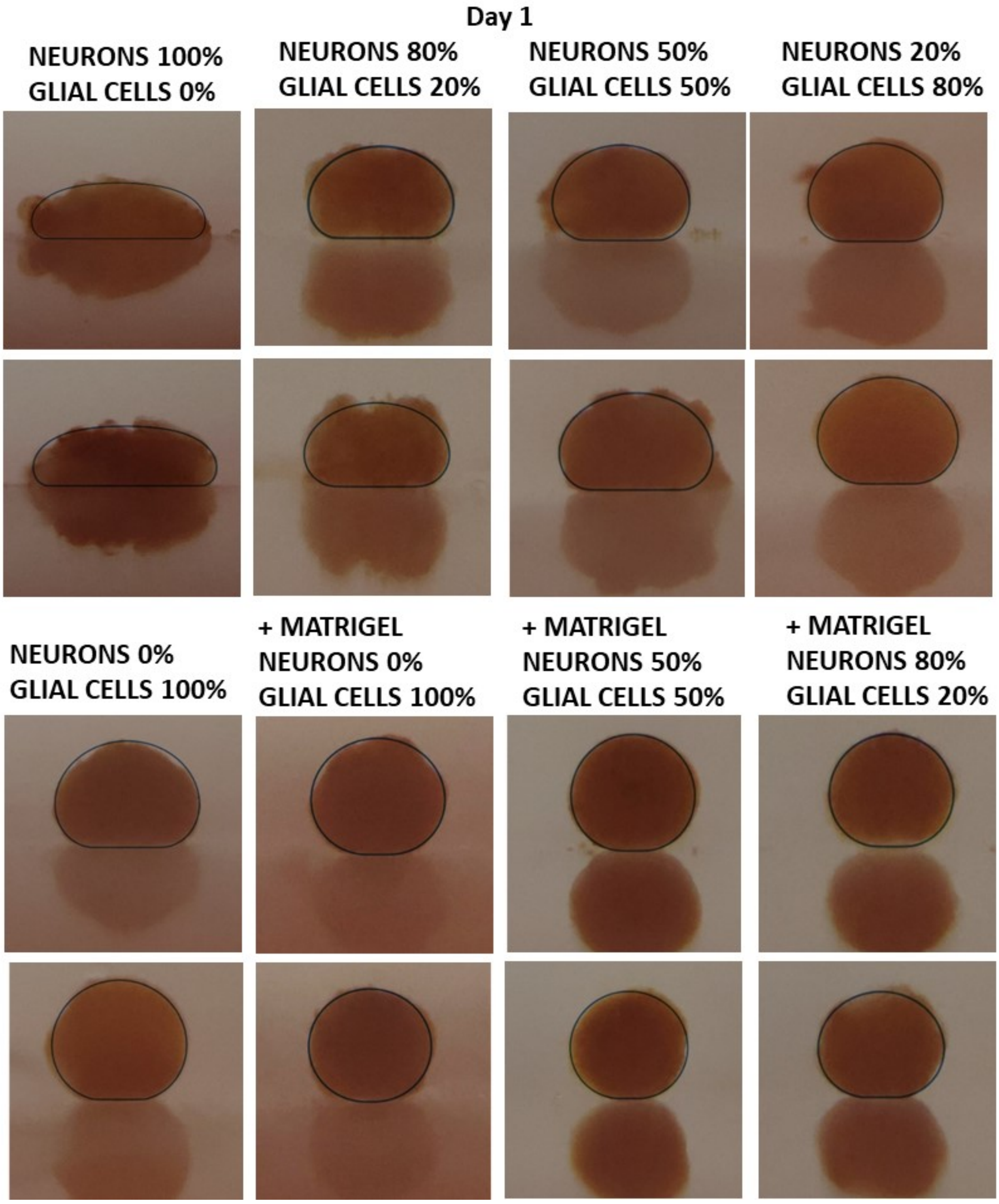


**Figure 10**. Spheroid lateral compression profiles for the different glial and neuronal intermixing ratio conditions at day 1 of maturation. The spheroids are compressed after 5 minutes of magnetic field gradient exposure.
